# Monoclonal antibodies for the S2 subunit of spike of SARS-CoV cross-react with the newly-emerged SARS-CoV-2

**DOI:** 10.1101/2020.03.06.980037

**Authors:** Zhiqiang Zheng, Vanessa M. Monteil, Sebastian Maurer-Stroh, Chow Wenn Yew, Carol Leong, Nur Khairiah Mohd-Ismail, Suganya Cheyyatraivendran Arularasu, Vincent Tak Kwong Chow, Raymond Lin Tzer Pin, Ali Mirazimi, Wanjin Hong, Yee-Joo Tan

**Affiliations:** Department of Microbiology and Immunology, Yong Loo Lin School of Medicine, National University Health System (NUHS), National University of Singapore.; Department of Laboratory Medicine, Karolinska Institute, Sweden; Public Health Agency of Sweden, Sweden; Bioinformatics Institute (BII), A*STAR (Agency for Science, Technology and Research), Singapore; Department of Biological Sciences (DBS), National University of Singapore.; National Public Health Laboratory (NPHL), National Centre for Infectious Diseases (NCID), Singapore; Institute of Molecular and Cell Biology (IMCB), A*STAR (Agency for Science, Technology and Research), Singapore; National Veterinary Institute, Sweden

**Keywords:** Coronavirus Disease 2019 (COVID-19), SARS-CoV-2, Spike protein, Cross-reactive antibodies

## Abstract

The emergence of a novel coronavirus, SARS-CoV-2, at the end of 2019 has resulted in widespread human infections across the globe. While genetically distinct from SARS-CoV, the etiological agent that caused an outbreak of severe acute respiratory syndrome (SARS) in 2003, both coronaviruses exhibit receptor binding domain (RBD) conservation and utilize the same host cell receptor, angiotensin-converting enzyme 2 (ACE2), for virus entry. Therefore, it will be important to test the cross-reactivity of antibodies that have been previously generated against the surface spike (S) glycoprotein of SARS-CoV in order to aid research on the newly emerged SARS-CoV-2. Here, we show that an immunogenic domain in the S2 subunit of SARS-CoV S is highly conserved in multiple strains of SARS-CoV-2. Consistently, four murine monoclonal antibodies (mAbs) raised against this immunogenic SARS-CoV fragment were able to recognise the S protein of SARS-CoV-2 expressed in a mammalian cell line. Importantly, one of them (mAb 1A9) was demonstrated to detect S in SARS-CoV-2-infected cells. To our knowledge, this is the first study showing that mAbs targeting the S2 domain of SARS-CoV can cross-react with SARS-CoV-2 and this observation is consistent with the high sequence conservation in the S2 subunit. These cross-reactive mAbs may serve as tools useful for SARS-CoV-2 research as well as for the development of diagnostic assays for its associated coronavirus disease COVID-19.

## Introduction

The Severe Acute Respiratory Syndrome Coronavirus (SARS-CoV) is the etiological agent for the infectious disease, SARS, which first emerged 17 years ago [1,2]. In December of 2019, another novel coronavirus (SARS-CoV-2) appeared to have crossed species barriers to infect humans and was effectively transmitted from person to person, leading to a pneumonia outbreak in Wuhan, China [3–5]. As of 1 March 2020, this coronavirus disease (COVID-19) has spread further within China and worldwide to 60 countries with over 87,000 confirmed cases and 2,980 fatalities [6]. The outbreak has been declared a Public Health Emergency of International Concern by the World Health Organization (WHO) and poses an increasing burden on global health and economy [6].

A study of 56 complete and partial SARS-CoV-2 genomes isolated from COVID-19 patients showed very high sequence conservation of more than 99%, indicating a recent introduction of the virus into the human population [7]. Phylogenetic analysis of SARS-CoV-2 revealed that like SARS-CoV and bat-derived SARS-like coronaviruses (SL-CoVs), it belongs to lineage B of the betacoronavirus genus [8,9]. While SARS-CoV-2 shares higher whole-genome sequence identity with bat-SL-CoVZC45 and bat-SL-CoVZXC21 (88 – 89%) than with SARS-CoV (79 – 82%), the receptor binding domain (RBD) of SARS-CoV-2 is more similar to SARS-CoV RBD, suggesting a possible common host cell receptor [8,9]. In line with this, several groups have demonstrated that SARS-CoV-2 utilizes the same host receptor, angiotensin-converting enzyme 2 (ACE2), as SARS-CoV for viral entry [3,10–12]. Due to its role in virus entry, the viral surface spike (S) glycoprotein has been the target for the generation of monoclonal antibodies (mAb). The S protein is functionally divided into two domains with the RBD-containing S1 subunit being responsible for attachment to host cell while the S2 subunit mediates fusion between viral and host membranes (reviewed by Li, F.) [13]. Although the origin of SARS-CoV-2 has not been established, SARS-CoV is believed to have originated from SL-CoVs residing in bats [14–17]. For the majority of SL-CoVs, the S1 subunit has low sequence homology to that of SARS-CoV which indicates species-dependent receptor binding [17,18]. On the other hand, the high amino acid sequence identity of more than 90% in the S2 subunit suggests that the fusion mechanism during virus infection is well-conserved [17,18].

In our previous work, we used an immunogenic fragment in the S2 subunit of SARS-CoV to generate a panel of murine mAbs [19].One of them, mAb 1A9, which binds to the S protein through a novel epitope within the S2 subunit at amino acids 1111-1130, has the ability to bind and cross-neutralize pseudotyped viruses expressing S of human SARS-CoV, civet SARS-CoV and bat SL-CoV strains [20]. In this study, we show that the sequence of the immunogen used to generate mAb 1A9, as well as three other mAbs, is conserved in multiple strains of SARS-CoV-2 and these mAbs bind to the spike protein of SARS-CoV-2 expressed in a mammalian cell line. Importantly, mAb 1A9 was demonstrated to detect S in SARS-CoV-2-infected cells.

## Materials and methods

### Cells

Vero E6 and COS-7 cells were purchased from American Type Culture Collection (Manassas, VA, USA) and cultured in Dulbecco’s Modified Eagle’s Medium (DMEM; Thermo Scientific) supplemented with 10% foetal bovine serum (FBS; HyClone) and penicillin-streptomycin solution (Thermo Fisher Scientific). Cells were maintained at 37 °C with 5% CO_2_.

### Purification of antibody

The hybridoma for mAb 1A9 was previously generated [19]. All mAbs were purified from cell culture supernatant using a HiTrap protein G HP affinity column (GE Healthcare) and stored at −80 °C. The purity of the mAb was confirmed by sodium dodecyl sulphate-polyacrylamide gel electrophoretic (SDS-PAGE) analysis. The concentration of the purified mAb was determined using the Coomassie Plus protein assay reagent (Thermo Scientific).

### Transient transfection and western blot analysis

COS-7 cells were seeded onto 6-well plates 24 hours prior to the transient transfection using Lipofectamine 2000 reagent (Thermo Scientific) according to the manufacturer’s protocol. SARS-CoV-2 S-expressing plasmids were codon-optimized and generated by gene synthesis (Bio Basic Asia Pacific) according to GenBank accession QHD43416.1. One plasmid is for expressing full-length S while the other is for expressing a fragment consisting of residues 1048-1206. At 48 hours post-transfection, cells were harvested, spun down by centrifugation and washed with cold PBS twice. Cells were then resuspended in RIPA buffer [50 mM Tris-HCl (pH 8.0), 150 mM NaCl, 0.5% NP40, 0.5% deoxycholic acid, 0.005% SDS and 1 mM phenylmethylsulfonyl fluoride] and subjected to five freeze-thaw cycles. Clarified supernatant containing the protein of interest was obtained by spinning down the cell lysate at 13,000 rpm at 4 °C to remove the cell debris and further analyzed by Western blot (WB) analysis. 20 μg of total cell lysates were resolved using electrophoresis on SDS-PAGE gels and transferred onto nitrocellulose membrane (Bio-Rad). The membrane was blocked in 5% skimmed milk in Tris-buffered saline with 0.05% Tween 20 (TBST) for 1 hour at room temperature and incubated with primary antibodies at 4 °C overnight. After the membrane was washed with TBST, it was incubated with a HRP-conjugated secondary antibody (Thermo Scientific) at room temperature for 1 hour. The membrane was then washed with TBST again and detected with enhanced chemiluminescence substrate (Thermo Scientific).

### Transient transfection and immunofluorescence analysis

For immunofluorescence (IF) analysis, COS-7 cells on glass coverslips were transfected as above and fixed at 24 hours post-transfection in 4% paraformaldehyde for 10 min at room temperature (RT) followed by permeabilization with 0.2% Triton X-100 (Sigma) for 5 min. Fixed cells were then blocked with PBS containing 10% fetal bovine serum for 30 min at RT. Cells were immunolabelled for 1 hour at RT with the indicated murine mAb and 45 min with Alexa Fluor 488-conjugated goat anti-mouse IgG antibody (Life Technologies). Immunolabelled coverslips were counterstained with DAPI (Sigma), and mounted using ProLong® Gold Antifade Mountant (Molecular Probes). Images were acquired with Olympus CKX53 microscope using Olympus LCAch N 20x/0.40 iPC objective lens and Olympus DP27 colour camera with Olympus cellSens software. Each channel was collected separately, with images at 1024 × 1024 pixels.

### Virus infection and IF

All works with live virus has been performed in the BSL3 facility at the Public Health Agency of Sweden. Vero-E6 cells were infected with SARS-CoV-2 (SARS-CoV-2-Iso/01/human/2020/SWE, GenBank accession no. MT093571) at a multiplicity of infection (MOI) of 1 in DMEM 2% FBS (Thermo Fisher). At 24h post-infection, cells were fixed with chilled methanol/acetone and the cells were kept in −20°C overnight. Cells were then stained using mAb 1A9 at 5μg/ml at 37°C for 30 minutes in immunofluorescence buffer (BSA 0.2%, Triton X100 0.1% in PBS, pH 7.4). They were then washed 3 times with PBS and incubated with goat anti-mouse conjugated to Alexa Fluor 488-conjugated goat anti-mouse IgG antibody (Thermo Fisher) containing DAPI in immunofluorescence buffer for an additional incubation of 30 minutes. Cells were washed 3 times with PBS before reading in fluorescent microscopy.

### Bioinformatics analysis

Spike glycoprotein reference protein sequences for SARS-CoV, SARS-CoV-2, batRaTG13, MERS and human common cold coronaviruses 229E, NL63, OC43 and HKU1 were downloaded from NCBI (GenBank accessions in alignment in Figure 1A). A multiple sequence alignment was created with MAFFT using the slow but accurate L-INS-I parameter settings [21] and the alignment curated, cut to the target region 1029-1192 (SARS-CoV numbering) and visualized with Jalview [22]. We used Mega X [23] to calculate the number of amino acid differences for all sequence pairs in the alignment of the mAb target region (Figure 1B) and the full S protein (Figure 1C) normalized by the length of the aligned sequence of the respective reference protein to obtain percent amino acid identities.

**Figure 1.**
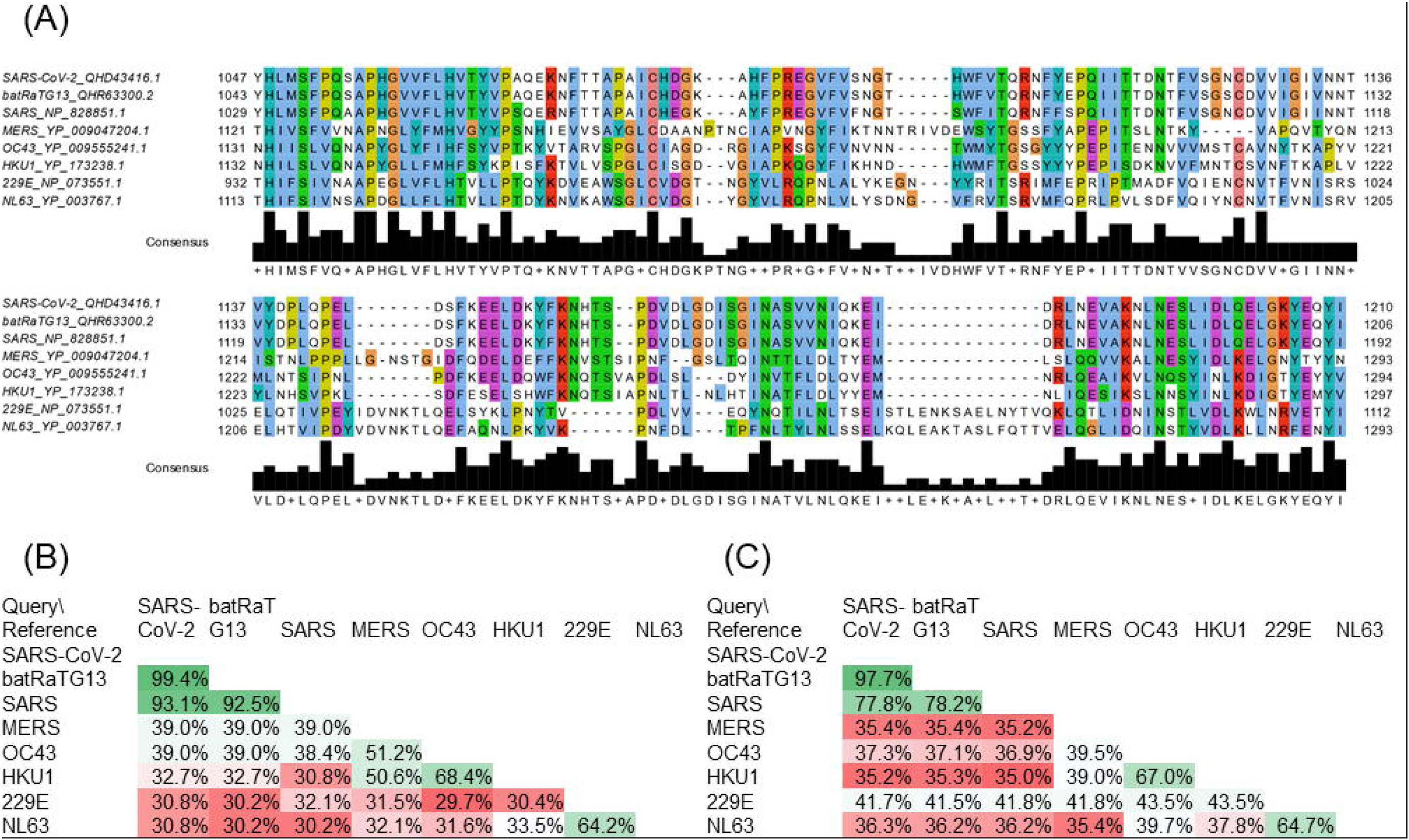
(A) Multiple sequence alignment across relevant coronaviruses for the fragment from S2 domain of SARS-CoV used to generate mAbs. (B) Pairwise amino acid identity (%) for fragment region. (C) Pairwise amino acid identity (%) for full spike protein.

To determine within-outbreak sequence diversity in the spike glycoprotein, 230 human and environmental outbreak virus sequences were downloaded from the GISAID database on March 1^st^ 2020. We gratefully acknowledge the authors, originating and submitting laboratories of the sequences from GISAID’s EpiCoV™ Database on which this part of the research is based. The list is detailed in Supplementary Table 1. The nucleotide sequences were searched with BLASTX against the reference spike glycoprotein. 174 hits covered the full length of the spike glycoprotein and amino acid mutations were counted and tabulated using a custom Perl script (Supplementary Table 2).

## Results

### An immunogenic domain in the S2 subunit of SARS-CoV is highly conserved in SARS-CoV-2 but not in endemic coronaviruses

By using five different fragments of SARS-CoV S to immunize rabbits, a fragment corresponding to residues 1029 to 1192 was found to stimulate neutralizing antibodies against SARS-CoV [24]. Interestingly, sequence alignment shows that this domain, which encompasses the heptad repeat (HR) 2 but not HR1, is highly conserved in SARS-CoV-2 (Figure 1A). When compared with additional reference sequences from bat RaTG13 (closest bat precursor), MERS and human common cold coronaviruses 229E, NL63, OC43 and HKU1 (Figure 1A), it becomes apparent that the amino acid identity between SARS-CoV-2 and SARS-CoV is much higher in this region (93%, Figure 1B) than over the full protein length (78%, Figure 1C) and the similarity drops sharply (<40% in this region) when considering MERS and the other coronaviruses infecting humans regularly.

We also studied within-outbreak sequence diversity across 174 full spike glycoproteins derived from nucleotide sequences shared via the GISAID platform (full acknowledgement in Supplementary Table 1) [25]. Only 4 amino acid mutations were found within the putative antibody binding region compared to 30 mutations over the full length protein (Supplementary Table 2). 2 of these 4 amino acid mutations are from a sequence flagged in GISAID’s EpiCoV^TM^ database as lower quality due to many undetermined bases.

### Four murine mAbs bind to a fragment of the S protein of SARS-CoV-2

Subsequently, a panel of murine mAbs was generated using SARS-CoV S(aa1029-1192) fragment and found to have neutralizing activities in vitro [19]. Four mAbs with distinct binding profiles, as mapped by internal deletion mutagenesis study, were selected for testing to determine if they cross-react with SARS-CoV-2. A fragment containing residues 1048 to 1206 of the S protein of SARS-CoV-2 was expressed in COS-7 cells via transient transfection and Western blot analysis was performed using four mAbs, namely 2B2, 1A9, 4B12 and 1G10. As shown in Figure 2A, all 4 mAbs detected this fragment of SARS-CoV-2, which is consistent with the sequence alignment shown in Figure 1A. Similarly, immunofluorescence analysis performed on transiently transfected COS-7 cells showed binding of the 4 mAbs to this S fragment of SARS-CoV-2 (Figure 2B).

**Figure 2.**
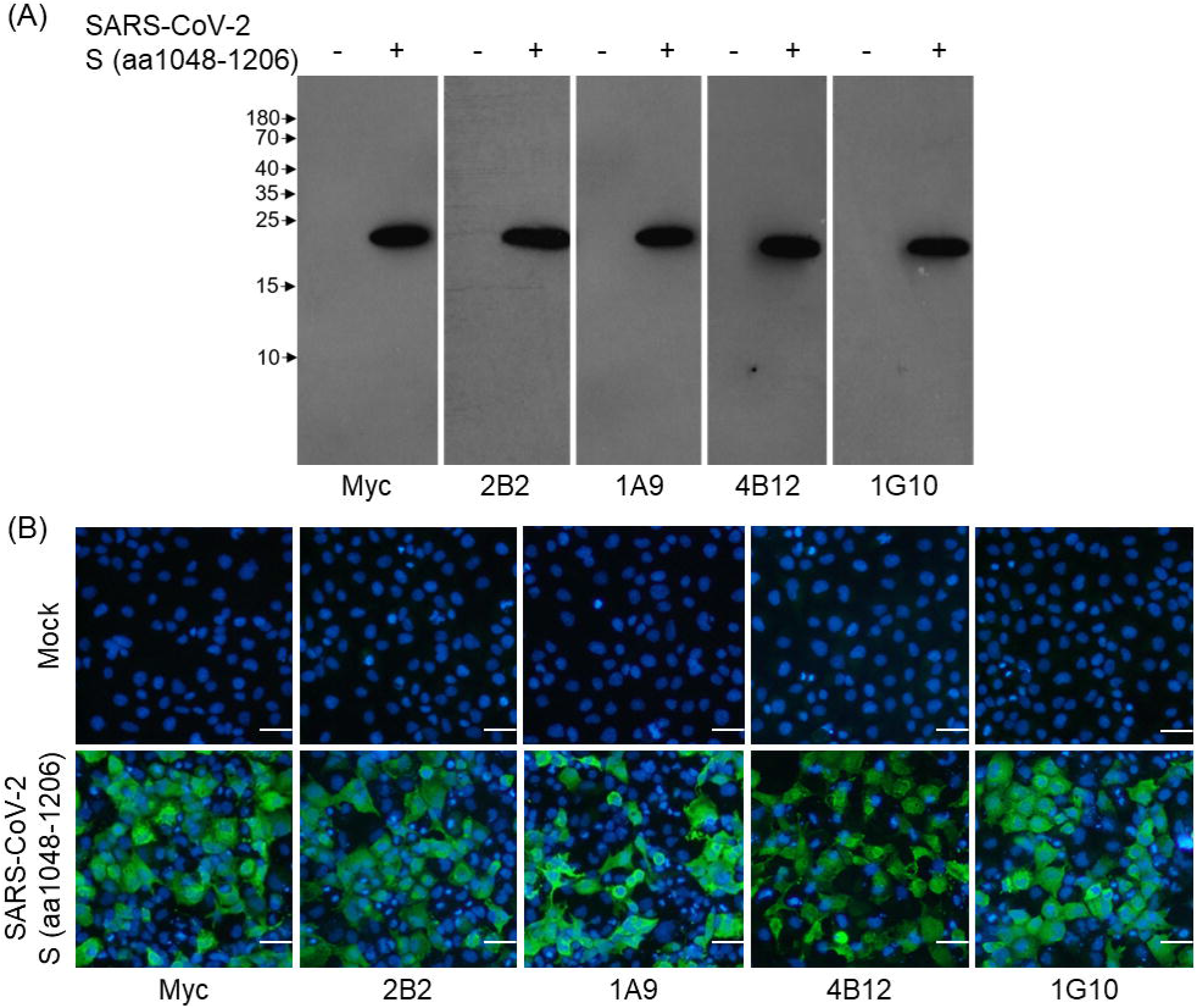
Mock-transfected COS-7 cells and cells expressing Myc-tagged SARS-CoV-2 S(aa1048-1206) fragment were used for (A) Western blot analysis using the indicated primary antibodies, followed by HRP-conjugated secondary antibody. (B) Immunofluorescence analysis using the indicated primary antibodies followed by Alexa Fluor 488-conjugated secondary antibody. Nuclei were counterstained with DAPI (blue). Scale bar = 50μm.

### Four murine mAbs bind to the full-length S protein of SARS-CoV-2

Next, the full-length S protein of SARS-CoV-2 was overexpressed in COS-7 cells and probed with each of the mAbs. As shown in Figure 3, all 4 mAbs bound to the full-length S protein of SARS-CoV-2.

**Figure 3.**
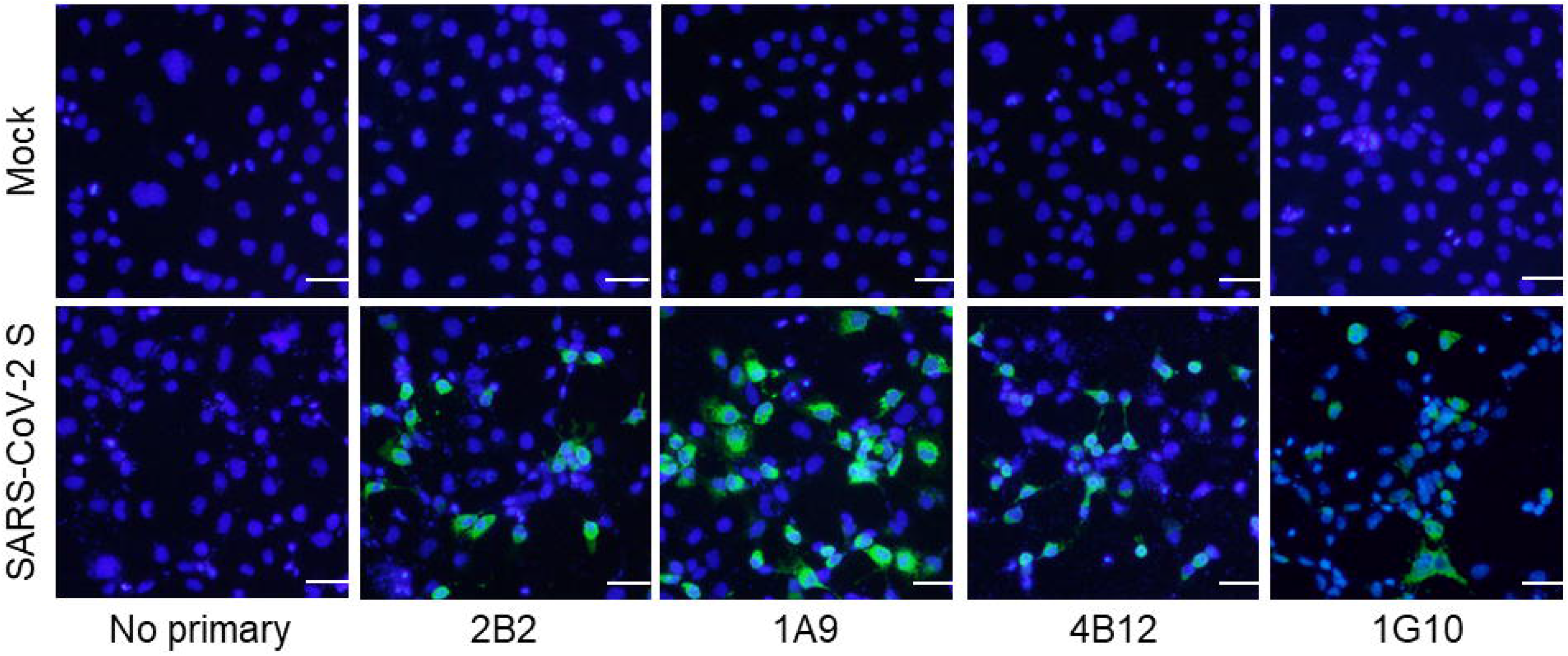
Immunofluorescence analysis was performed on mock-transfected COS-7 cells and cells expressing full-length SARS-CoV-2 S protein. The indicated primary antibodies were used followed by Alexa Fluor 488-conjugated secondary antibody. Nuclei were counterstained with DAPI (blue). Scale bar = 50μm.

### Mab1A9 binds to S expressed in SARS-CoV-2-infected cells

Since mAb 1A9 was previously shown to have broad cross-reactivity to civet SARS-CoV and bat SL-CoV strains [20], it was tested on SARS-CoV-2-infected Vero-E6 cells. As shown in Figure 4, mAb 1A9 stained a significant number of SARS-CoV-2-infected cells at 24h post-infection showing that it is sensitive enough to detect the expression of S during infection.

**Figure 4.**
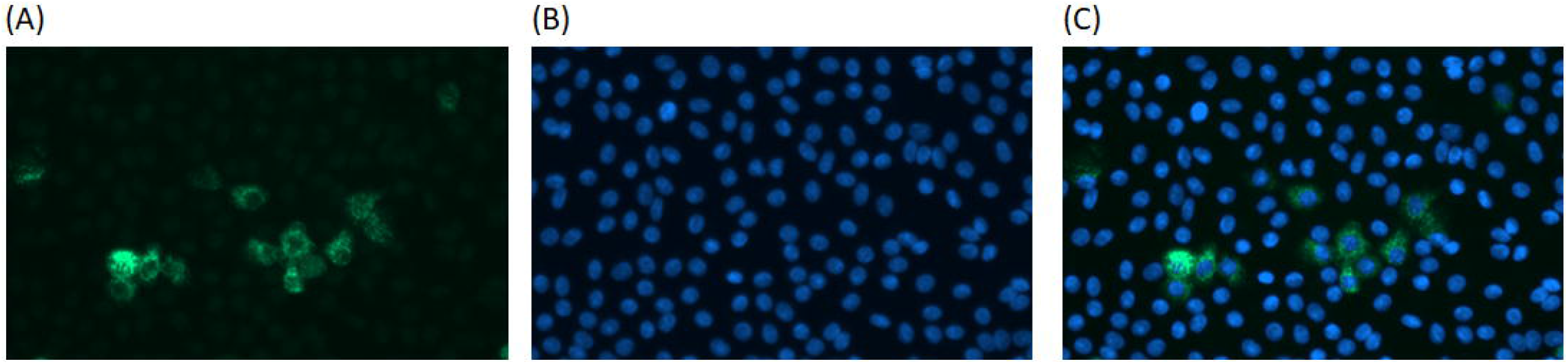
(A) Vero E6 cells were mock-infected or infected with SARS-CoV-2 (MOI of 1). At 24h post infection, the cells were stained with mAb 1A9 (5μg/ml) followed by Alexa Fluor 488-conjugated secondary antibody. (B) Nuclei were counterstained with DAPI (blue). (C) Merged image

## Discussion

Numerous mAbs against the S protein of SARS-CoV have been generated for research and diagnostic assay development. Some of these may be able to cross-react with the S protein of SARS-CoV-2 and serve as tools to aid research on this newly emerged virus. In this current study, an immunogenic domain in the S2 subunit of SARS-CoV was found to be highly conserved in multiple strains of SARS-CoV-2. Consistently, WB and IF analyses showed that 4 different mAbs generated using this SARS-CoV domain were cross-reactive against the S protein of SARS-CoV-2. In addition, mAb 1A9 stained a significant number of SARS-CoV-2-infected cells at 24h post-infection showing that it is sensitive enough to detect the expression of S during infection. Thus, these mAbs will be useful for studying the kinetics of SARS-CoV-2 replication in vitro and development of diagnostic assays for COVID-19. It is noteworthy that cytotoxic T-lymphocyte (CTL) epitopes also reside at residues 884-891 and 1116-1123 within the S2 subunit of SARS-CoV [26]. Interestingly, the latter CTL epitope overlaps with the epitope recognized by mAb 1A9 [20]. Hence, the S2 subunit may serve as an important antigen for inducing both humoral as well as cell-mediated immunity against SARS-CoV and SARS-CoV-2.

Recent cross-reactivity studies have evaluated SARS-CoV neutralizing antibodies that bind to the RBD-containing S1 subunit. Although both SARS-CoV and SARS-CoV-2 use ACE2 as a receptor for viral entry [3,10], several SARS-CoV RBD-directed mAbs did not cross-react with SARS-CoV-2 RBD [27,28]. Interestingly, CR3022, which was isolated from a SARS convalescent patient, showed cross-reactivity to SARS-CoV-2 RBD and recognizes an epitope that does not overlap with the ACE2 binding site [28]. To our knowledge, this is the first study showing that mAbs targeting the S2 domain of SARS-CoV can cross-react with SARS-CoV-2 and this observation is consistent with the high sequence conservation in the S2 subunit. Besides the mAbs characterized here, several other mAbs have been reported to bind to epitopes in the S2 subunit of SARS-CoV [29–31]. Thus, it will be important to determine if these mAbs can also cross-react with SARS-CoV-2.

## Acknowledgements

The work performed in NUS/NUHS was supported by NUHS Research Office under Project Number NUHSRO/2020/033/RO5+5/CORONAVIRUS/LOA (WBS R-571-000-071-733). The work performed in IMCB and BII was also supported by A*STAR through intramural funding and an A*CRUSE gap funding (ACCL/19-GAP064-R20H-F).

